# Low-temperature HILIC provides enhanced separations and stability for LC-MS-based metabolomics

**DOI:** 10.64898/2025.12.19.695573

**Authors:** Yifan Liu, Madison L. Jastrab, Michael Xiao, Miriam Lisci, Taysir K. Bader, Alexis A. Jourdain, Thomas E. Wales, Owen S. Skinner

## Abstract

Liquid chromatography–mass spectrometry is a potent and robust tool for studying metabolism. However, conventional workflows can suffer from poor peak shapes, limited pressure tolerance, co-elution of polar metabolites, and unstable retention times. Here, we describe the development of a more stable HILIC method for LC-MS metabolomics of human plasma and cell extracts, optimizing a zwitterionic HILIC (Z-HILIC) column for improved untargeted performance. We found that using high-pH ammonium bicarbonate with 90% acetonitrile in mobile phase B (ABC B) can greatly improve peak shapes of select metabolites when compared to 100% acetonitrile (ACN B), but at the cost of poor retention time stability. We therefore focused on optimizing chromatography for the ACN B method and observed that cooling the column to 5 °C substantially enhanced peak shape. This low-temperature Z-HILIC (LT-ZHILIC) method provides high-resolution separation of metabolites from both cellular extracts and human plasma, is stable over days, and generally outperformed a standard method using the widely described ZIC-pHILIC column. Application of the untargeted LT-ZHILIC method to characterize the metabolic consequences of glutamine and pyruvate deficiency in human cells revealed a striking change in nucleotide phosphates; a perturbation that was not observed in the ZIC-pHILIC analysis of the same samples likely due to inadequate elution profiles. In sum, the LT-ZHILIC workflow offers a robust platform to advance untargeted metabolomics by improving metabolite coverage, resolution, and retention time stability, making it a promising technique for providing novel insights into cellular metabolic rewiring and the human plasma metabolome.

## Introduction

Untargeted metabolomics provides a comprehensive and unbiased view of metabolism by profiling a wide range of known and unknown metabolites, making it a beneficial tool to investigate metabolic perturbations and pathway alterations across diverse biological and environmental systems.^1–3^ This discovery-oriented approach enables hypothesis generation and the identification of novel biomarkers and pathways, including early metabolic changes in cardiovascular and metabolic diseases. Application of untargeted metabolomics has highlighted the role of the “dark metabolome” in biomarker discovery, and uncovered alterations in bile acid, amino acid, and glycan metabolism.^3,4^ Of particular interest are polar metabolites like sugars, amino acids, nucleotides, and organic acids, which play central roles in energy metabolism and signaling, and whose altered abundances often reflect disease states, drug responses, or nutritional status.^5^ However, the chemical diversity, low abundance, and instability of these metabolites make their reliable detection challenging, underscoring the need for robust LC–MS methodologies.

Moreover, complex biological samples such as plasma, tissues, and cell extract often generate overlapping signals, adducts, in-source fragments, and isomeric species, all of which further complicate accurate metabolite identification.^6^

Among analytical platforms, LC–MS has proven highly effective due to its superior sensitivity, mass accuracy, and broad metabolite coverage.^7^ However, the choice of chromatographic column is critical for accurate metabolite identification. Reverse-phase LC (RP-LC) columns can generate narrow and well-defined chromatographic peak shapes, ^8^ but very polar and charged compounds are often poorly retained, eluting in an unresolved “void volume” at the start of the chromatogram.^9^ This lack of retention hinders separation and increases the likelihood of co-elution, especially in complex biological mixtures where metabolites may share similar physicochemical properties, including isomeric forms. ^10^ To extend beyond these challenges, mixed-mode columns which combine reversed-phase and ion-exchange or polar functionalities have been developed to broaden metabolite coverage within a single run.^11^ These columns can improve selectivity for compounds with a wide range of polarities, though their dual retention mechanisms often require careful optimization to ensure reproducible separations.

To overcome the limitations of RP-LC for highly polar metabolites, hydrophilic interaction liquid chromatography (HILIC) has emerged as a core strategy in metabolomics. HILIC employs a polar stationary phase and high-organic mobile phase, enabling strong retention of charged and hydrophilic compounds such as amino acids, nucleotides, and sugar phosphates that are poorly resolved by RP-LC.^12^ This mode of separation is particularly effective for untargeted metabolomics, where comprehensive coverage of central carbon and energy metabolism is critical. Several stationary phase chemistries have been developed, including bare silica, amino, amide and zwitterionic (sulfobetaine) ligands, each providing distinct selectivity through a combination of liquid partitioning, hydrogen bonding, and electrostatic interactions.^13,14^ Among zwitterionic HILIC columns, ZIC-pHILIC is one of the most widely used in untargeted metabolomics. It features a polymer (“p”) linkage that bonds the sulfobetaine stationary phase to a polymeric support, providing strong retention of highly polar metabolites and broad metabolome coverage, which has made it a standard platform for polar metabolite profiling. However, the polymeric nature of the stationary phase also introduces important limitations. In addition to lower backpressure tolerance (<200 bar) and reduced mechanical robustness compared with hybrid or silica-based materials, polymer supports cannot be manufactured in the smaller particle sizes characteristic of UPLC/UHPLC technologies. This constraint limits achievable peak capacity and diminishes the potential gains in sensitivity and resolution, ultimately affecting long-term stability and reproducibility in large-scale metabolomics analyses.^15^ To address these weaknesses, one newer design has introduced bridged ethylene hybrid (BEH) particles functionalized with zwitterionic ligands, giving rise to the “Z-HILIC” column.^16^ This design provides enhanced mechanical strength and substantially greater pressure tolerance compared with conventional ZIC-pHILIC, enabling more robust operation under UHPLC conditions. Separately, both Z-HILIC and ZIC-pHILIC benefit from being packed into inert column hardware, which reduces undesirable metal interactions and protects metal-sensitive analytes. Although the mechanisms differ Z-HILIC uses inert metallic hardware while ZIC-pHILIC employs PEEK-lined material both approaches help minimize analyte losses associated with metal surfaces.^17^ Direct evaluation of the two columns demonstrated that Z-HILIC achieves sharper peak shapes, improved resolution of isomeric species, and greater metabolite coverage than ZIC-pHILIC under similar conditions. ^18^ However, despite its high promise, certain metabolites remain poorly resolved or exhibit suboptimal peak shapes on Z-HILIC, and the long-term stability of the column under routine use conditions remains insufficiently characterized.

Two main chromatographic approaches have been established for Z-HILIC separations, each offering distinct advantages and compromises. The first method uses 15 mM ammonium bicarbonate at high pH in both mobile phases (ABC B), with mobile phase B consisting of 15 mM ammonium bicarbonate in 90% acetonitrile. This approach provides high efficiency and strong retention of polar analytes particularly for metabolites in central carbon metabolism.^19^ In contrast, the second approach uses high-pH ammonium bicarbonate only in mobile phase A with pure acetonitrile as mobile phase B (ACN B), improving the separation and detection of highly polar intermediates and broadening overall metabolite coverage. ^20^ Despite their widespread use, the relative stability and long-term performance of these two methods have not been systematically evaluated.

Here, we compare both methods and find that the ABC B method, while generally providing better peak shapes than the ACN B method for a broad range of metabolites, exhibits pronounced retention time drift after just one day. Although this instability is often overlooked in the metabolomics literature, it poses a significant limitation for large-scale studies that require reproducibility and long-term stability. To address these challenges, we optimized the ACN B method to enhance peak shapes while preserving its superior retention time stability. Through these efforts, we have developed a low-temperature Z-HILIC (LT-ZHILIC) strategy that yielded substantial improvements in peak quality and led to improved chromatographic resolution for metabolites from both human plasma and intracellular extracts. Unlike conventional methods that rely on elevated temperatures to improve peak shapes and reduce tailing, our approach operates at 5 °C, yielding improved separation efficiency while maintaining reproducibility across a wide range of metabolites, including those from cell extracts, plasma, and complex cultured-cell metabolomes.

Notably, in untargeted metabolomics of nutrient-deprived cultured cells, untargeted computational analysis detected key intracellular nucleotide phosphates such as ATP, ADP, IMP, and GMP with the LT-ZHILIC method, but not ZIC-pHILIC, highlighting that a robust chromatographic method is essential to capture biologically relevant metabolic changes under stress. Collectively, our results establish LT-ZHILIC as a robust, high-performance alternative that overcomes the limitations of existing Z-HILIC workflows by combining retention stability with superior chromatographic resolution.

## Experimental section

### Chemical and LC-MS reagents

For chromatographic separation, two gradient systems were prepared. The first (ABC B) comprised mobile phase A (15 mM ammonium bicarbonate in MS-grade water; Fisher Scientific) and mobile phase B (15 mM ammonium bicarbonate in 90% acetonitrile; Fisher Scientific), with the pH of mobile phase A adjusted to 9.2 using MS-grade ammonium hydroxide (Fisher Scientific). The second (ACN B) system used mobile phase A (20 mM ammonium bicarbonate in MS-grade water; Fisher Scientific) and mobile phase B (100% acetonitrile; Fisher Scientific), with the pH of mobile phase A likewise adjusted to 9.2 using MS-grade ammonium hydroxide (Fisher Scientific). The LT-ZHILIC method was initiated at 0.15 mL/min with 80% mobile phase B at 5 °C from 0.0 to 0.5 min. From 0.5 to 20.5 min, the gradient was held at 20% mobile phase B at the same flow rate, and maintained until 21.3 min. The column was then equilibrated at 0.15 mL/min and 80% mobile phase B from 21.5 to 26.0 min, followed by a brief flushing step at 0.3 mL/min and 80% mobile phase B from 26.0 to 26.1 min.

### Preparation of cell and plasma extractions

K562 cells (purchased from ATCC) were reauthenticated by STR profiling in 2025. Cells were cultured in Dulbecco’s Modified Eagle Medium (DMEM; Fisher) supplemented with 10% fetal bovine serum (FBS; Fisher) under standard conditions (37 °C, 5% CO₂). For metabolite extraction, three biological replicates of K562 cells were prepared at a density of 1 × 10⁶ cells/mL in a 6-well plate. Each sample was transferred to a microcentrifuge tube and centrifuged at 400 × g for 5 min at room temperature, after which the supernatant was carefully aspirated to remove the culture medium. Intracellular metabolites were extracted by adding 600 µL solvent mixture (40% methanol, 40% acetonitrile, 20% LC–MS grade water, 240 µL /240 µL /120 µL) containing 0.1 M formic acid. Samples were vortexed briefly and incubated on ice for 10 min. To neutralize the extract, 42 µL of 150 mg/mL ammonium bicarbonate was added to each sample. This extraction strategy was modified from a previously described method. ^21^

For plasma metabolite extraction, a mixture containing 40% methanol (MS grade; Fisher), 40% acetonitrile (MS grade; Fisher), and 20% pooled human plasma (Fisher) was prepared to a final volume of 600 µL, vortexed for 10 seconds, and incubated on ice for 10 min.

Both cell and plasma extracts were centrifuged at 21000 × *g* for 10 min at 4 °C. The resulting supernatants were transferred to LC vials and analyzed under varying chromatographic conditions, including ammonium bicarbonate (ABC) and acetonitrile (ACN) mobile phases on different HILIC columns.

### Untargeted metabolomics of glutamine and pyruvate supplemented K562 cells

K562 cells (ATCC) were cultured in Dulbecco’s Modified Eagle Medium (DMEM; Gibco, A14430-01) supplemented with 25 mM glucose (Sigma, G7021), 2 mM L-glutamine (Amimed, 5-10K00-H), 2 mM sodium pyruvate (Invitrogen, 11360039), 10% fetal bovine serum (FBS; Fisher), and 100 U/mL penicillin–streptomycin (Bioconcept). Cells were maintained under standard conditions (37 °C, 5% CO₂) in T25 or T75 flasks and subjected to four treatment conditions: (+Gln/+Pyr), (+Gln/−Pyr), (−Gln/+Pyr), and (−Gln/−Pyr). For nutrient-deprivation conditions, water was substituted for the omitted component. K562 cells were seeded at 0.5 × 10⁶ cells/mL for +glutamine conditions (12.5 × 10⁶ cells in 25 mL) or 0.6 × 10⁶ cells/mL for −glutamine conditions (15 × 10⁶ cells in 25 mL) and cultured in T75 flasks at 37 °C with 5% CO₂ overnight. The following day, 20 × 10⁶ cells from each condition were collected, resuspended in 40 mL of freshly prepared medium, and split into four T25 flasks (10 mL each) as technical replicates. After 5 h of incubation, cells were harvested by centrifugation at 300 × *g* for 3 min, washed once with PBS, and flash-frozen in liquid nitrogen for metabolite extraction. The extraction procedure was same as non-treated K562 cells mentioned previously.

### Instrumentation (LC-MS analysis and cryogenic LT-ZHILIC setup)

Samples were maintained at 4 °C in the autosampler before analysis. LC–MS experiments were performed on a Vanquish UHPLC system coupled to an Orbitrap Exploris 480 mass spectrometer (Thermo Fisher Scientific). Chromatographic separations were carried out using a Waters Premier BEH Z-HILIC column (1.7 µm, 2.1 × 150 mm; item no. 186009983) equipped with a matching VanGuard pre-column (1.7 µm, 2.1 × 5 mm; item no. 186009984). For comparison, the same LC–MS setup was used with a ZIC-pHILIC column (2.1 × 150 mm; Sigma-Aldrich, item no. 1.50460.0001). Column temperature was varied according to method conditions to evaluate chromatographic performance.

Modified chromatographic conditions are described in Table 1. The original ABC B LC method was operated at a flow rate of 0.30 mL/min, with the column temperature set to 30 °C and the initial mobile phase B composition at 90%. The original ACN B method was run at a flow rate of 0.15 mL min⁻¹, with the column temperature maintained at 30 °C and the initial mobile-phase B composition at 80%. The optimized LT-ZHILIC method was operated at a flow rate of 0.15 mL min⁻¹, column temperature of 5 °C, and a starting mobile-phase B composition of 80%. The mass spectrometer was operated at a resolution of 600,000 with a scan range of m/z 70–1000 in polarity-switching mode (positive and negative). The H-ESI ion source was used with static spray voltages of +3.2 kV (positive) and –2.8 kV (negative). Gas settings were fixed: sheath gas = 35, auxiliary gas = 5, sweep gas = 1, ion-transfer tube temperature = 320 °C, and vaporizer temperature = 175 °C

**Table 1.** Summary of key method parameters for chromatographic optimization. Flow rate, column temperature, and initial percentage of solvent B, from the starting ACN B for method **M0** to **M9**, with results shown in **Figure 3**.

| Method name | Flowrate (ml/min) | Temperature (°C) | Initial B % |
| --- | --- | --- | --- |
| M0 | 0.15 | 30 | 80 |
| M1 | 0.3 | 30 | 80 |
| M2 | 0.15 | 30 | 95 |
| M3 | 0.15 | 50 | 80 |
| M4 | 0.15 | 10 | 80 |
| M5 | 0.1 | 10 | 80 |
| M6 | 0.15 | 10 | 85 |
| M7 | 0.15 | 10 | 90 |
| M8 | 0.15 | 10 | 95 |
| M9 | 0.15 | 5 | 80 |

For MS/MS fragmentation, positive and negative modes were acquired separately using the AcquireX Deep Scan workflow. In negative mode, the intensity threshold was set to 5 × 10⁴, with dynamic exclusion enabled after one occurrence, an exclusion duration of 15s, and a mass tolerance of 10 ppm.

Cryogenic temperature experiments were conducted by enclosing the Z-HILIC column in a custom insulated foam housing connected to an external refrigerated chamber. ^22^ This setup was required because the UHPLC column compartment could not be operated below 4 °C. The chamber was maintained at various sub-zero temperatures (< 0 °C) and directly coupled to the LC–MS system to ensure stable low-temperature operation.

### Data processing and metabolites identification (freestyle and compound discoverer)

Metabolite profiles from each replicate were analyzed using Freestyle (Thermo Fisher Scientific) to evaluate peak shapes. Four representative metabolites: (iso)citrate, malate, ATP, and glutamate were selected for peak-shape comparison, and their chromatograms were extracted directly from Freestyle. Retention time comparisons were performed in Skyline (University of Washington). A transition list for key metabolites, including 2-ketoglutarate, glutamate, and succinate, was manually generated based on their precursor m/z values, using [M – H]⁻ as the precursor ion for negative-mode detection. Retention time data were exported from Skyline and analyzed in Prism (GraphPad) to calculate mean values and standard deviations.^1^

For comprehensive metabolite identification, cultured-cell extracts were analyzed using both ZIC-pHILIC and LT-ZHILIC methods, followed by pooled-sample analysis with an AcquireX deep-scan workflow in Xcalibur (Thermo Fisher Scientific) to obtain MS/MS spectra. The resulting data were processed in Compound Discoverer (Thermo Fisher Scientific) using the following filters: matched MS/MS spectra, non-blank compound names, correct reference-ion polarity (+ for positive mode, – for negative mode), peak rating ≥ 7, and mzCloud best-match score ≥ 50. Compound identification employed a four-tiered library search strategy encompassing mzCloud, predicted compound matches, mass-list searches, ChemSpider, and Metabolika databases. Volcano plots were generated in Prism for statistical and comparative visualization of differential metabolite features.

## Results

### Comparing the ABC B and ACN B gradient systems

To evaluate the performance of the Z-HILIC column for untargeted metabolomics, we applied the ABC B and ACN B methods to two representative biological matrices: intracellular metabolites isolated from K562 human myelogenous leukemia cells and circulating metabolites from human plasma. We selected (iso)citrate, malate, and ATP (K562) or glutamate (plasma), which exhibited distinct elution profiles that allowed us to directly evaluate the performance of each method **(Figure 1)**. In cell extracts, (iso)citrate showed comparable elution profiles between the ABC B method (orange traces) and the ACN B method (purple traces) **(Figure 1A)**, whereas malate and ATP displayed sharper and more symmetrical peaks with the ABC B method **(Figures 1B-C)**. Similar improvements in peak shape were observed for other metabolites in plasma samples **(Figures 1D–F)**. These results indicate that the ABC B method produced sharper elution profiles for the selected metabolites in both plasma and intracellular extracts. Notably, malate in K562 cells showed the largest improvement, with the full peak width decreasing from 2.28 ± 0.08 min under the ACN B method to 1.21 ±0.42 min using ABC B **(Figure 1B)**.

**Figure 1.**
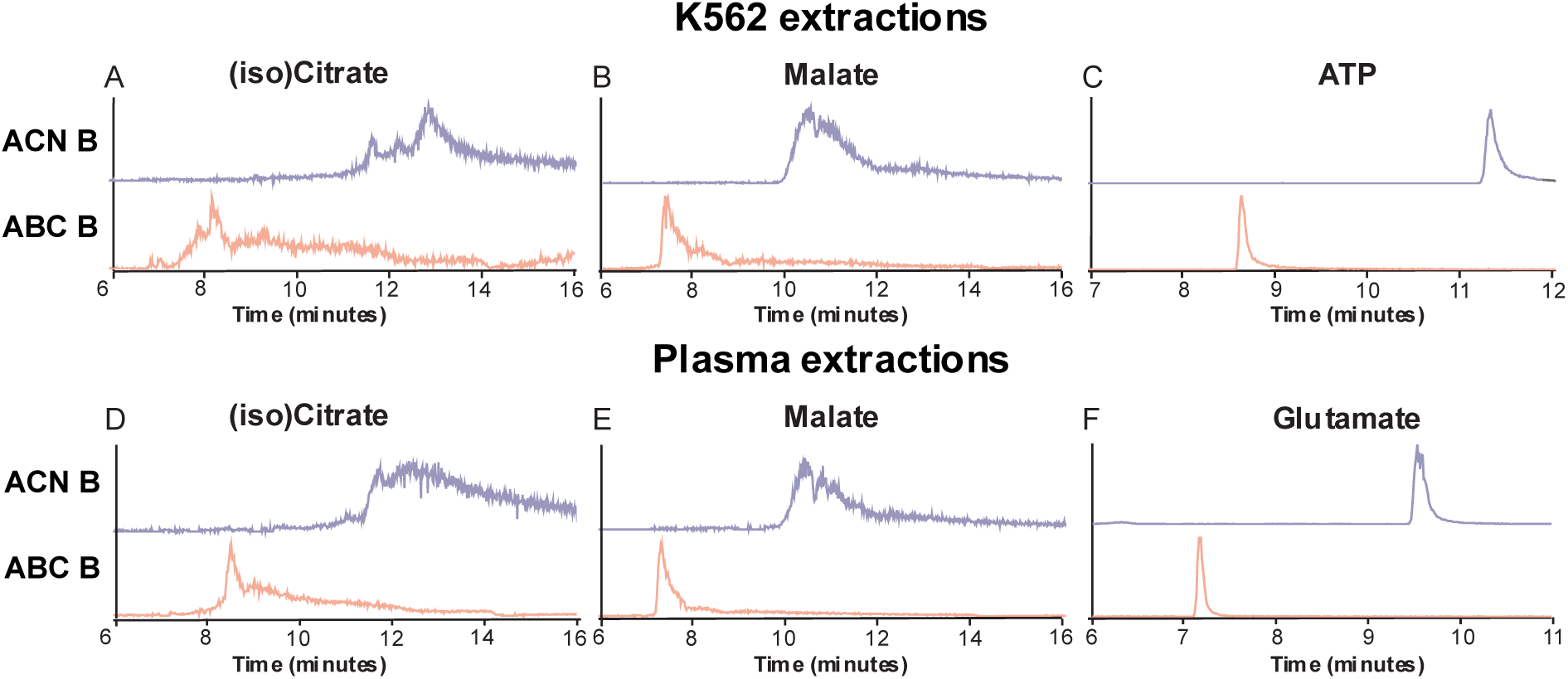
Metabolite peak shape comparison between ACN B and ABC. **B.** (A–F) Representative chromatograms of three selected metabolites, (iso)citrate, malate, and aspartate (n=2 replicates for ABC B, n=3 replicates for ACN B) in K562 cell extracts (A–C) and plasma samples (D–F) run using the ACN B and ABC B methods. Column details are mentioned in experimental section.

We next wanted to characterize the retention time stability of each method across multiple days, a critical characteristic for the successful analysis of large batches of samples that is common practice for metabolomics workflows. We conducted a four-day time-course experiment to evaluate the stability of the ABC B and ACN B mobile phase systems. We evaluated glutamate, alpha-ketoglutarate, and succinate due to their consistently well-resolved elution profiles with both separation in both cell extracts **(Figure 2A–B; Supplementary Figure S1A)** and plasma extracts **(Figure 2C–D; Supplementary Figure S1B)** to monitor retention time precision. With ABC B (orange traces), the retention time of the 2-ketoglutarate peak from cell extracts progressively shifted from 6.79± 0.01 minutes to 6.30 ± 0.01 minutes from day 0 to day 3 **(Figure 2A)**. The corresponding ΔRT plots further emphasized this observation: ABC B exhibited an average retention time drift of ∼0.5 minutes by day 3 relative to day 0 of all 3 replicates of 2-ketoglutarate, whereas ACN B showed minimal deviation (<0.01 minutes) across the four-day period **(Figures 2E-F)**. Similar patterns were observed for glutamate in both cell and plasma extracts **(Figures 2C–D, G-H)**. Collectively, our results demonstrate that while ABC B results in improved peak shapes, it suffers from severe retention time drift that doesn’t affect ACN B, suggesting that ACN B has better performance for larger sample batches.

**Figure 2.**
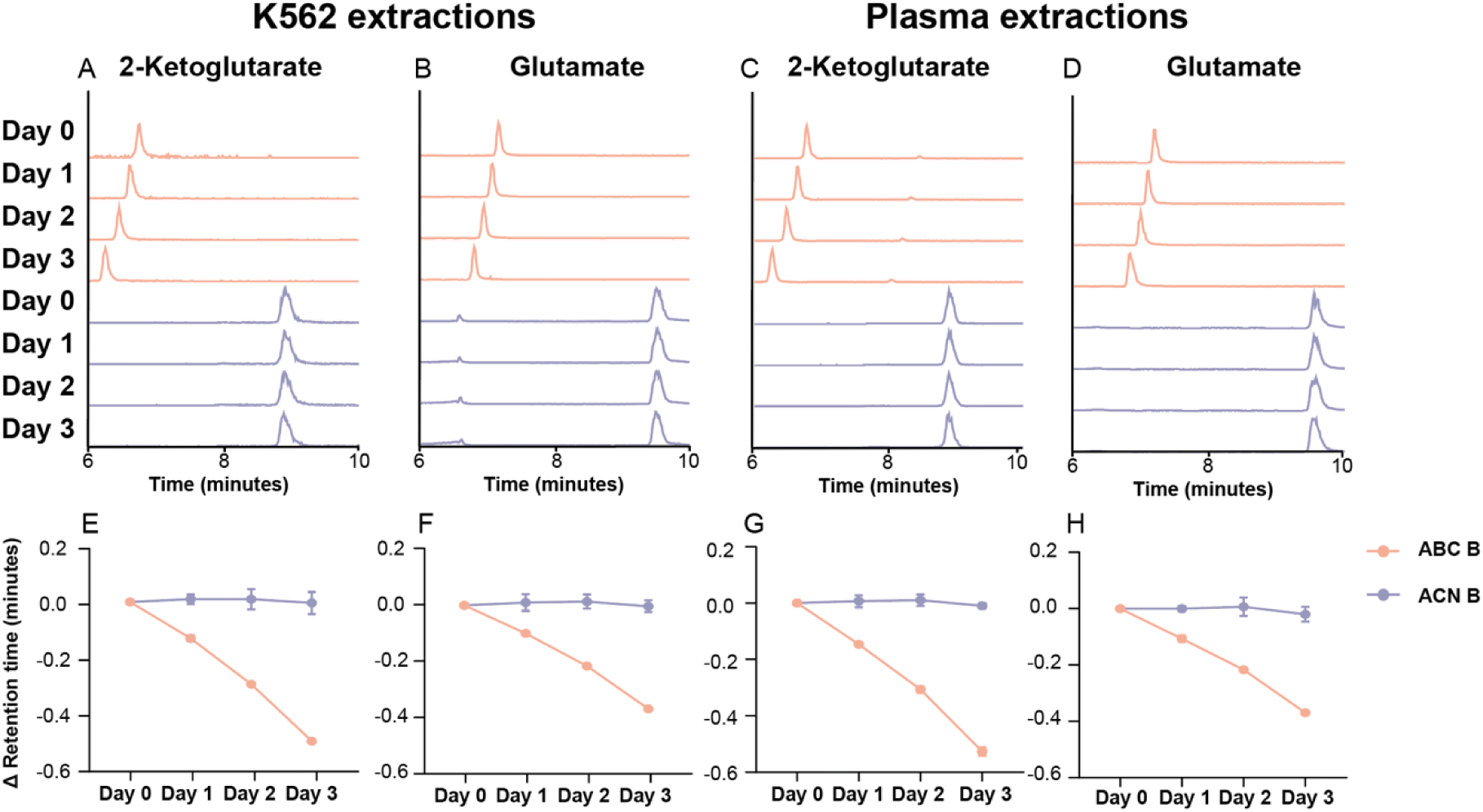
Retention time stability of ACN B and ABC. **B.** (A–D) Representative chromatograms of 2-ketoglutarate and glutamate in K562 cell extracts (A–B) and plasma samples (C–D). Metabolites analyzed using the ABC B method are shown in purple; those analyzed using the ACN B method are shown in orange. (E–H) retention time trends for each metabolite over a three-day time course (Day 0 to Day 3) comparing the ABC B (purple) and ACN B (orange) methods.

### Optimization of ACN B chromatography

The ACN B method maintained stable retention times but yielded suboptimal peak shapes, primarily due to significant tailing, most notably for (iso)citrate and malate **(Figure 1)**. To systematically address these limitations, we optimized chromatographic conditions by varying flow rate, column temperature, and the initial percentage of mobile phase B, generating a series of methods described in **Table 1 (M0–M9)**. The original ACN B protocol **(M0, Figures 1,2)**, which uses 80% initial mobile phase B, a flow rate of 0.15 mL/min, and a column temperature of 30 °C, **M0**, served as the baseline condition. We used the (iso)citrate, malate, and ATP or glutamate from intracellular and plasma extracts **(as in Figure 1)** to assess changes in peak quality. Increasing the flow rate to 0.3 mL/min **(M1)** yielded little improvement; (iso)citrate peaks remained jagged and poorly resolved **(Figure 3A)**. Similarly, raising the initial mobile phase B to 95% (**M2**) did not improve peak symmetry and introduced unrelated background peaks. Consistent with prior reports^7^, column temperature adjustments had more pronounced effects. Increasing the temperature to 50 °C **(M3)** improved peak quality for ATP in cell extracts and glutamate in plasma extracts (**Figures 3C, 3F**), though (iso)citrate and malate continued to display broad, tailing peaks. Decreasing the temperature to 10 °C **(M4)** improved chromatographic performance across these metabolites, yielding more symmetrical peaks. Building on this observation, we tested additional refinements of flow rate and mobile phase composition **(M5–M8)** under low-temperature conditions but observed only modest benefits, as (iso)citrate and malate peaks remained broad and jagged. The final optimized method **(M9)**, employing 80% mobile phase B, a flow rate of 0.15 mL/min, and a column temperature of 5 °C, delivered the best overall targeted analysis. This method, which we termed “low-temperature Z-HILIC” or LT-ZHILIC, produced consistently narrow peak widths and improved peak symmetry for all tested metabolites, with notable improvements in (iso)citrate and malate peak shapes **(Figures 3A–B, 3D–E; light green traces)**. Collectively, our results demonstrate that lowering the column temperature is an effective strategy for improving peak shape in the ACN B method, establishing LT-ZHILIC as a robust method for metabolomics applications.

**Figure 3.**
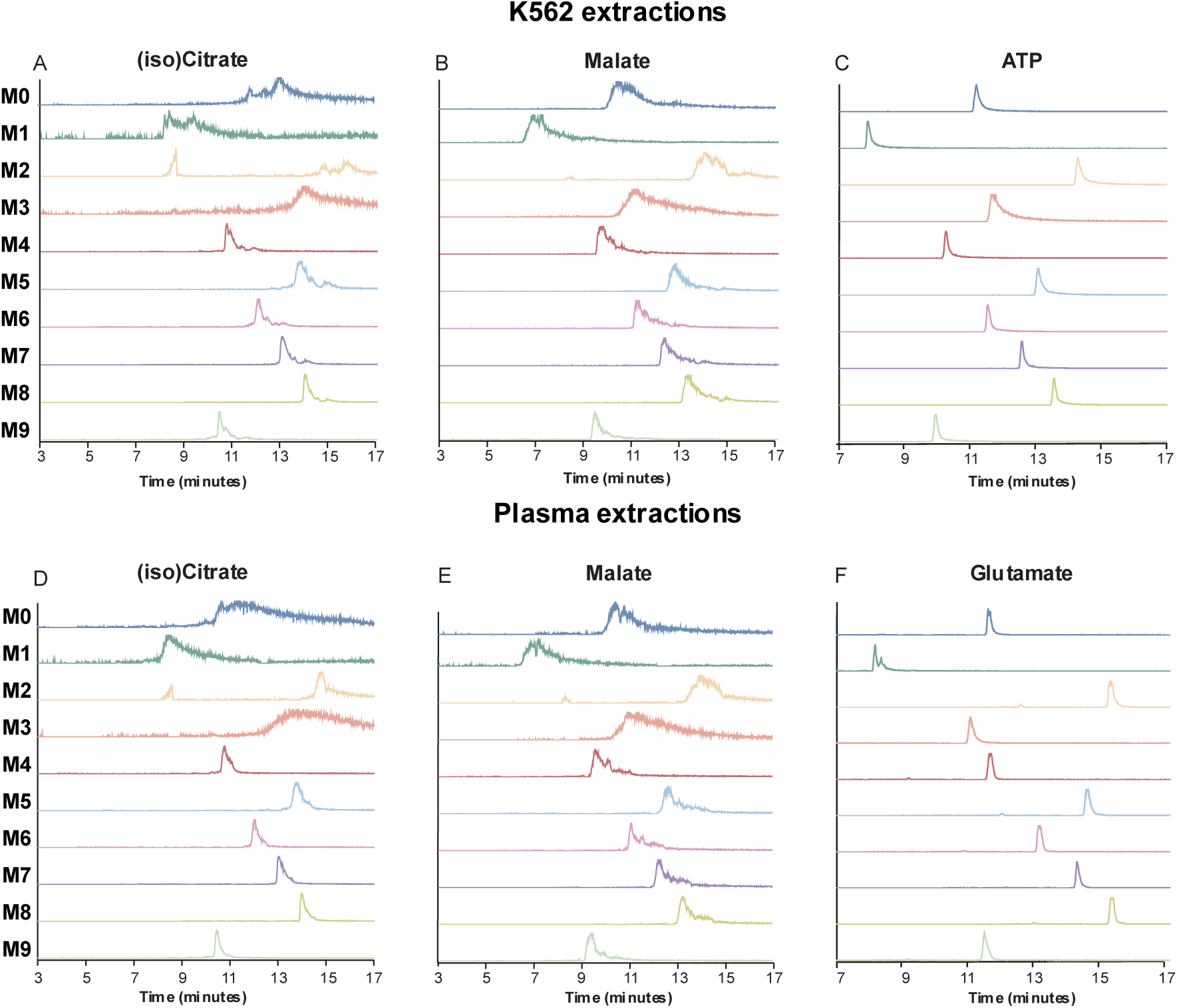
Optimization of the ACN B method using the method parameters (M0-M9) described in. Table 1. (A–C) Representative chromatograms of (iso)citrate, malate, and ATP in K562 cell extracts; (D–F) chromatograms of (iso)citrate, malate, and glutamate in plasma samples. M0 represents the baseline ACN B method, using a flow rate of 0.15 mL/min, a column temperature of 30 °C, and an initial composition of 80% mobile phase B. The parameters for each subsequent optimization step are summarized in Table 1. The final optimized method (M9), which employed a flow rate of 0.15 mL/min, a column temperature of 5 °C, and an initial 80% B, yielded the best overall peak performance.

Building on the low-temperature results, we next examined whether further cooling could enhance chromatographic separation. To this end, we maintained the Z-HILIC column under cryogenic conditions, with temperatures lowered to –20 °C using a custom-designed temperature-controlled chamber to assess performance stability and peak resolution **(Supplementary Figure S2A–D)**. We analyzed the same representative metabolites from both intracellular and plasma extracts for direct peak-shape comparison **(Supplementary Figure S3, A–F)**. Relative to the LT-ZHILIC method at 5 °C (light green traces), further reductions in temperature (–5 °C, –10 °C, –20 °C; darker green traces, top to bottom) did not yield improvements. In fact, at –10 °C, peak shapes deteriorated noticeably, for example (iso)citrate in cell extractions **(Supplementary Figure S3 A)**. These findings indicate that the optimal performance is achieved with the LT-ZHILIC method at 5 °C, while lower cryogenic temperatures compromise chromatographic quality.

To confirm that the LT-ZHILIC method exhibits the same retention time stability as ACN B, we next assessed the LT-ZHILIC method in a four-day time-course experiment. As in the previous analysis **(Figure 2)**, we selected and monitored 2-ketoglutarate and glutamate **(Figure 4A–D)**. Chromatograms of 2-ketoglutarate from cell extracts across days 0 to 3 revealed little retention time drift, and the corresponding ΔRT plot confirmed deviations of less than 0.023 minutes **(Figure 4A)**. We observed similar stability for other metabolites in both cell and plasma extracts, with ΔRT values remaining negligible throughout the four-day period **(Figures 4B–D; Supplementary Figures S4A–B)**. Collectively, our results demonstrate that LT-ZHILIC enhances chromatographic performance while providing stable retention times over long analytical sequences. This stability is critical for large-scale metabolomics, where minimizing retention drift ensures consistent feature alignment, preserves metabolite detectability, and enables reliable comparison across hundreds of samples.

**Figure 4.**
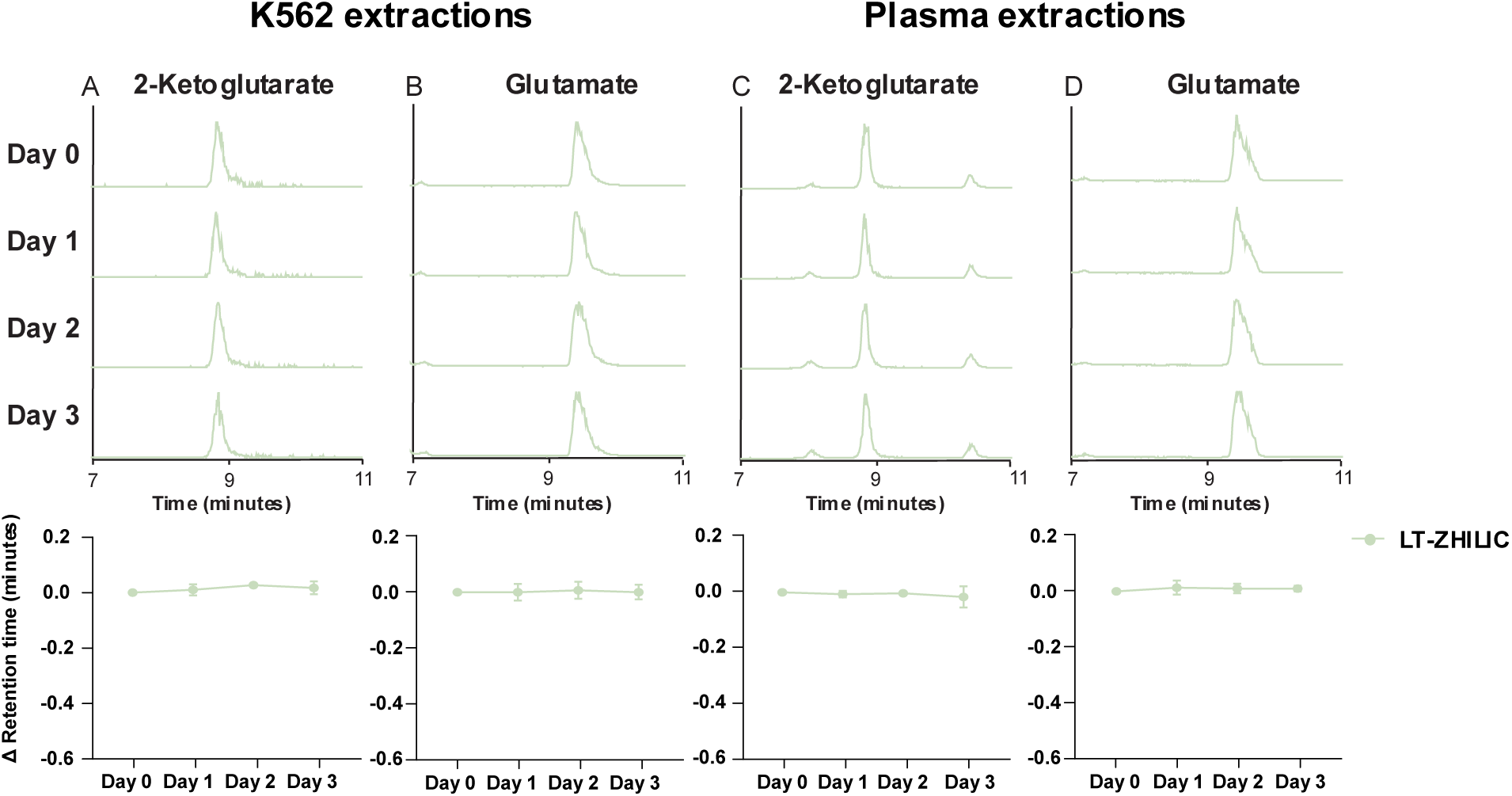
Retention time stability of LT-ZHILIC. Time-course analysis of LT-ZHILIC stability in K562 and plasma sample extracts. (A–B) 2-ketoglutarate and glutamate from K562 extracts. (C–D) 2-ketoglutarate and glutamate from plasma extracts. Over the four days, LT-ZHILIC demonstrated consistent stability.

### Comparison of ZIC-pHILIC and LT-ZHILIC performance

We next compared the LT-ZHILIC method to an established method on the ZIC-pHILIC column.^20,23^ In cell extracts, (iso)citrate exhibited comparable peak symmetry and sharpness between ZIC-pHILIC (blue trace) and LT-ZHILIC (light green trace) (**Figure 5A**). Similarly, other metabolites showed consistent peak symmetry and sharpness across both columns in cell and plasma extracts **(Figures 5B–F)**. Notably, ATP exhibited sharper and narrower peaks under LT-ZHILIC conditions, highlighting its improved chromatographic performance. These findings indicate that LT-ZHILIC achieves peak shapes equivalent to, and in some cases surpassing, that of the widely used ZIC-pHILIC column, underscoring its potential as a robust alternative for metabolomics applications.

**Figure 5.**
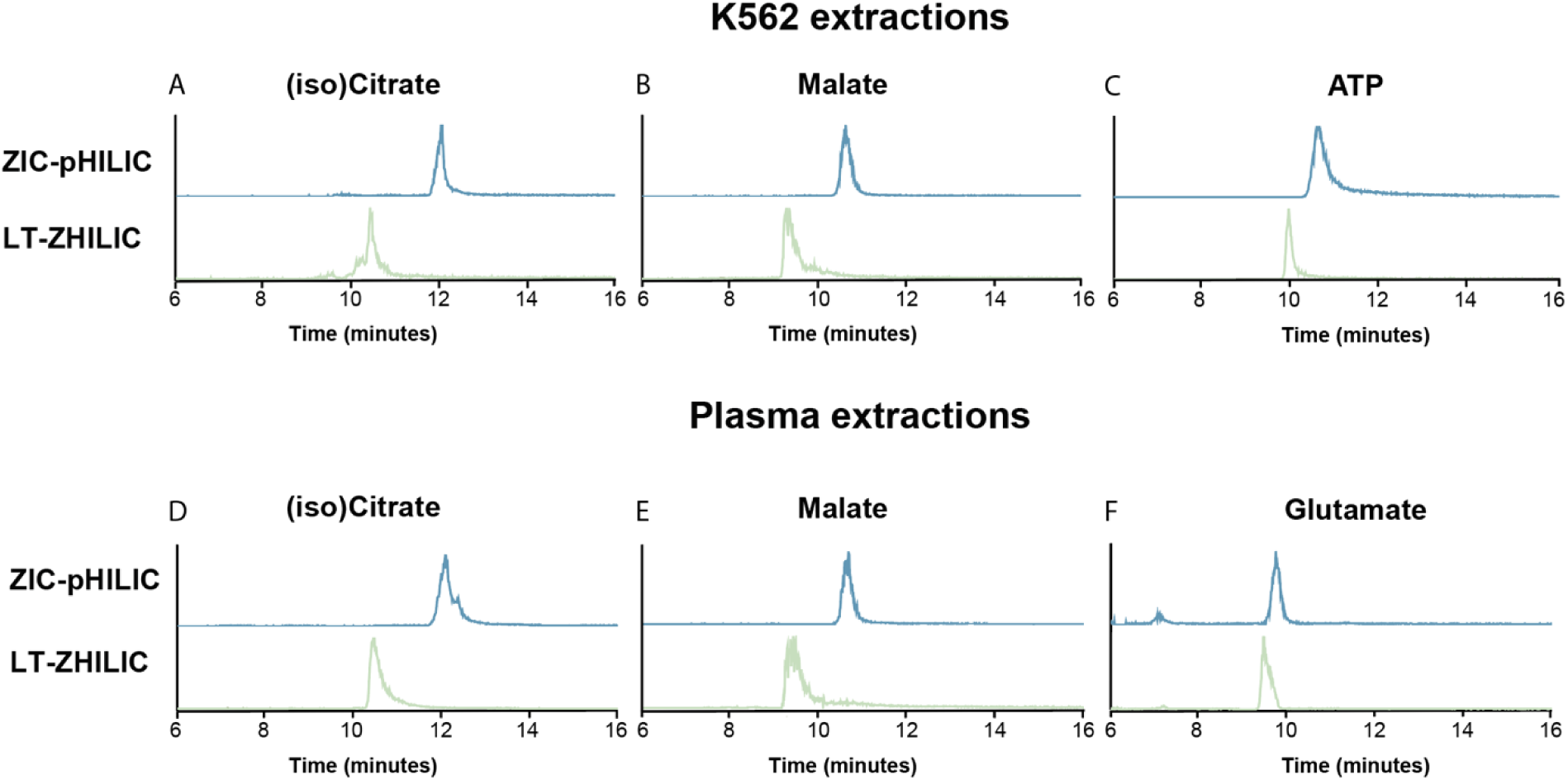
Peak shape comparison between LT-ZHILIC and ZIC-pHILIC. Peak-shape comparison of selected metabolites analyzed on ZIC-pHILIC and LT-ZHILIC (A-F). (A-C) representative results from K562 cell extracts and (D–F) corresponding results from plasma extracts. Traces from ZIC-pHILIC are shown in blue, and those from Z-HILIC in light green.

Given the peak shape improvement with using the Z-HILIC column at low temperature, we next applied the low-temperature strategy to ZIC-pHILIC at 5 °C to test whether performance could be further improved **(Supplementary Figure S5A–F)**. In K562 extracts, (iso)citrate showed no improvement compared to the original ZIC-pHILIC method at 30 °C (blue traces); the 5 °C ZIC-pHILIC runs (pink traces) even produced less symmetrical peaks with more tailing **(Supplementary Figure S5A)**. Other metabolites similarly showed no enhancement with the low-temperature approach. For instance, glutamate displayed an unrelated small peak eluting at an earlier retention time in the 5 °C ZIC-pHILIC method (**Supplementary Figure S5F)**. Overall, these results indicate that the benefits of the low-temperature strategy do not translate to ZIC-pHILIC. We therefore used ZIC-PHILIC at 30°C for the further performance comparisons.

### Targeted metabolomics analysis of cultured cell extracts using LT-ZHILIC

To further evaluate the performance of the LT-ZHILIC method in complex biological samples, we cultured K562 cells under four nutrient conditions varying in glutamine and pyruvate availability: (1) complete medium containing both substrates (control), (2) glutamine only, (3) pyruvate only, and (4) medium lacking both glutamine and pyruvate. We selected (Iso)citrate, malate, and ATP as representative metabolites for targeted analysis. While malate and ATP exhibited comparable peak areas between the two methods **(Figures 6A–C)**, (iso)citrate showed markedly enhanced signal intensity under LT-ZHILIC conditions, indicating improved detection sensitivity. Variability assessment further underscored the advantage of LT-ZHILIC: in 10 of 12 treatment–metabolite comparisons, LT-ZHILIC yielded lower coefficients of variation (CV%) than ZIC-pHILIC **(Figures 6D–F)**. Although we observed slightly higher CVs under specific conditions (e.g., glutamine-enriched but pyruvate-deprived media), LT-ZHILIC overall demonstrated comparable or superior intensity and generally higher ability to reliably quantify these metabolites, supporting its high suitability for targeted metabolomics workflows.

**Figure 6.**
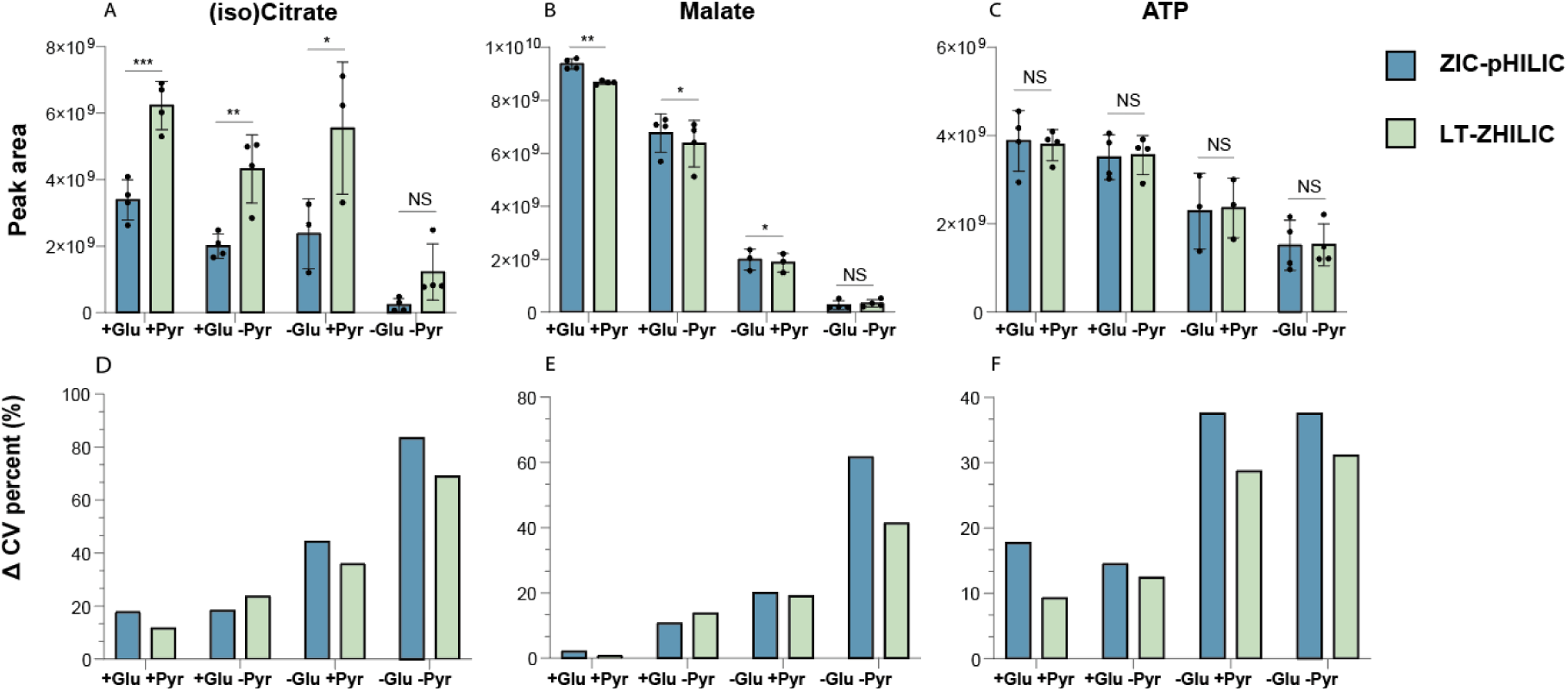
**LT-ZHILIC vs. ZIC-pHILIC quantitation of metabolites in cells with nutrient deficiency**. (A-F) (A–C) Comparison of peak areas across four treatment conditions analyzed on ZIC-pHILIC (blue) and LT-ZHILIC (green), comparing the sensitivity of both methods for treated biological samples (D–F) Coefficient of variation (CV%) analysis of both methods for each group.

To further probe the targeted metabolomics analysis capabilities of LT-ZHILIC, we examined the NADH and NAD⁺ profiles under each nutrient condition. These cofactors were chosen because they play central roles in maintaining cellular redox balance and serve as sensitive indicators of oxidative stress during nutrient perturbation. NADH, NAD⁺, and their redox ratios were quantified by both ZIC-pHILIC and LT-ZHILIC across four treatment groups **(Figure 7A–C)**. NADH level measurements were higher with ZIC-pHILIC compared to LT-ZHILIC across all treatments **(Figure 7A).** We observed a similar decrease for NAD⁺ in LT-ZHILIC, although the magnitude of the change was smaller **(Figure 7B).**

**Figure 7.**
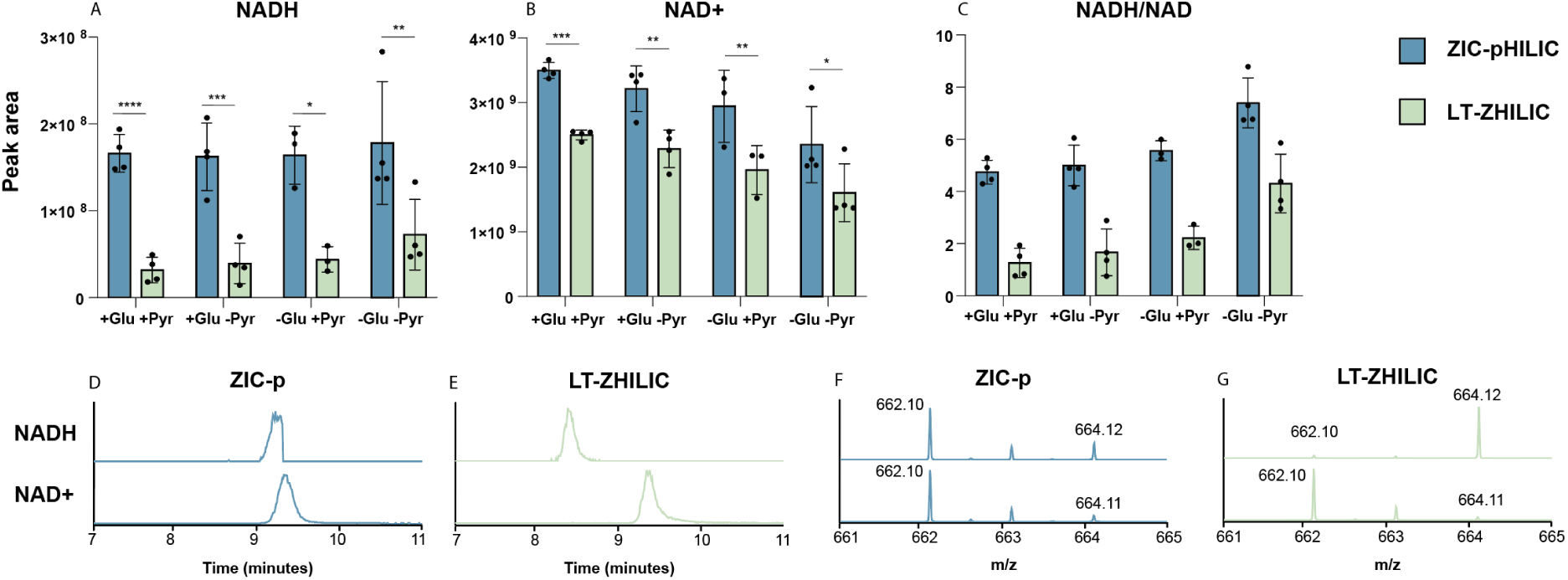
Targeted redox quantitation of redox of NADH and NAD⁺ redox cofactors. (A–C) Peak areas of NADH and NAD⁺ in K562 cell extracts analyzed using ZIC-pHILIC (blue) and LT-ZHILIC (green). NADH and NAD⁺ co-elute on ZIC-pHILIC, whereas LT-ZHILIC achieves clear baseline separation (D–F). Corresponding mass spectra illustrating overlapping signals in ZIC-pHILIC compared to distinct spectral resolution in LT-ZHILIC (E-G).

Correspondingly, the measured NADH/NAD⁺ redox ratio was also substantially higher in ZIC-pHILIC than in LT-ZHILIC **(Figure 7C).** To investigate the underlying reason for these discrepancies, we directly compared chromatographic and spectral profiles of NADH and NAD⁺ between the two methods. Because NADH and NAD⁺ share nearly identical molecular weight and polarity, their separation is challenging for most chromatographic systems. In ZIC-pHILIC, NADH and NAD⁺ co-eluted, producing overlapping chromatographic peaks **(Figure 7D)** and nearly indistinguishable mass spectra **(Figure 7F).** The fragment data shows the M+2 isotopic peak of NAD⁺ overlapped with the M+0 peak of NADH, leading to artificially elevated NADH quantification and poor peak shape. In contrast, LT-ZHILIC achieved clear baseline separation of these species, yielding distinct chromatographic peaks **(Figure 7E)** that lead to well-resolved mass spectra **(Figure 7F).** These results demonstrate the ability of LT-ZHILIC to resolve structurally related redox cofactors such as NADH and NAD⁺. Together with the consistently higher ratios observed in ZIC-pHILIC, this suggests that ZIC-pHILIC may overestimate NADH and NAD⁺ abundance and ratio, potentially leading to misleading interpretations of cellular redox state and associated pathway alterations.

### Untargeted metabolomics analysis of nutrient deprivation using LT-ZHILIC and ZIC-pHILIC

To investigate the ability of both methods to characterize metabolic alterations in cultured cells and uncover pathway-level changes, we applied untargeted metabolomics to analyze the four nutrient conditions described above. We measured the same samples in full scan polarity-switching mode using both ZIC-pHILIC and LT-ZHILIC and used a Compound Discoverer automated pipeline to analyze metabolite abundances in each condition. We generated volcano plots comparing each nutrient-deprived condition compared with control conditions **(Figure 8A-F, Supplementary Figure S6A-F).** Overall, LT-ZHILIC identified 148 metabolites, compared with 138 detected using ZIC-pHILIC through the same pipeline. Across most treatment conditions in negative-ion mode, LT-ZHILIC detected a greater number of significantly altered metabolites (p < 0.01, FC > 2) when glutamine and/or pyruvate were removed relative to the full-nutrient control (+Gln/+Pyr), although in some cases both methods yielded comparable numbers of significant changes. For example, in negative ion mode, combined glutamine/pyruvate deprivation yielded only 8 upregulated metabolites with ZIC-pHILIC **(Figure 8C)**, whereas LT-ZHILIC detected 19 significantly altered metabolites under the same condition **(Figure 8F)**. Similar trends were observed across other treatments as well as positive mode (**Figures 8A, B, D, E and Supplementary Figure S6A-F**), where LT-ZHILIC consistently detected more significant metabolite changes than ZIC-pHILIC. **(Figures 8G–H and Supplementary Figure S6G–H).** The exact mechanism underlying these differences between the two columns remains unclear. It is possible that LT-ZHILIC provides improved fragmentation or more consistent elution profiles, which, when analyzed with the same metabolomics pipeline, resulted in a higher number of detected metabolites.

**Figure 8.**
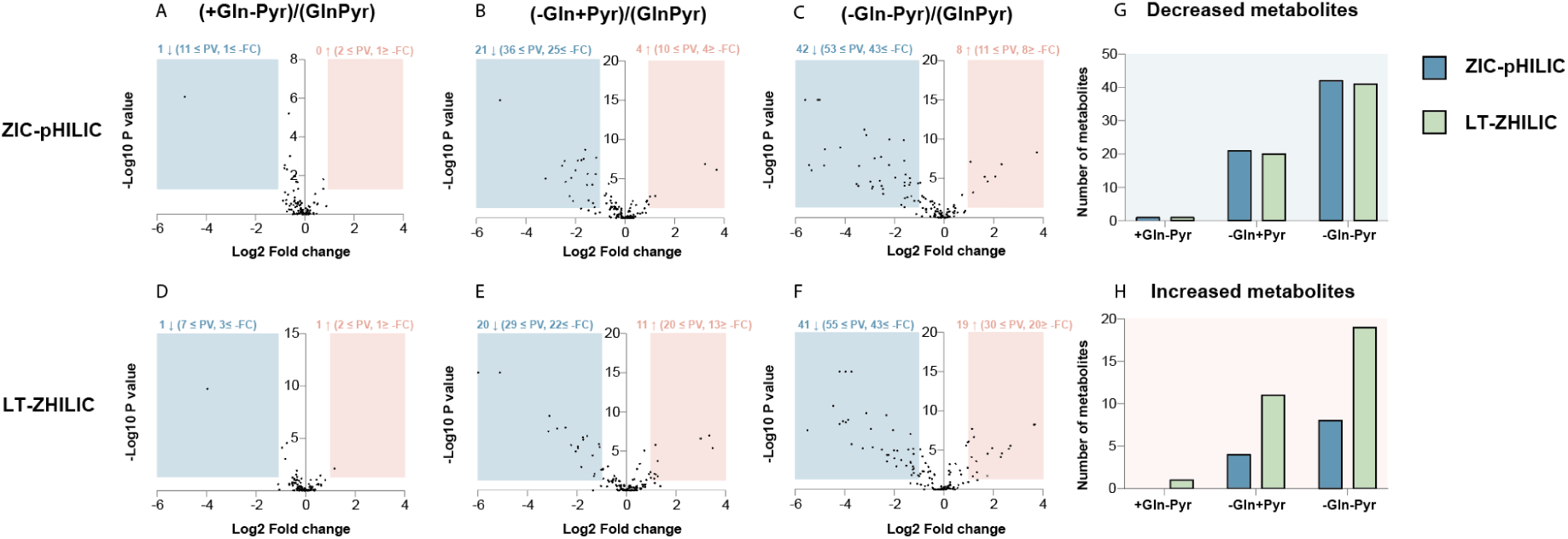
Negative-mode volcano plots of K562 cell extracts under different treatment conditions with or without glutamine and pyruvate. Analyzed with a Compound Discoverer automated pipeline from data acquired using ZIC-pHILIC (A-C). Corresponding results analyzed using LT-ZHILIC (D-F). (G–H) Summary of downregulated (G) and upregulated (H) metabolites detected using ZIC-pHILIC (blue) and LT-ZHILIC (green).

We next performed a focused analysis of the significantly altered metabolites in negative ion mode under combined glutamine and pyruvate deprivation. Several nucleotide phosphate species—including TTP, ATP, ADP, IMP, and GMP—showed significant alterations only when analyzed using LT-ZHILIC (highlighted in red), whereas they were not identified in ZIC-pHILIC analyses **(Figure 9A–B)**. To explore the source of this improved performance, we compared the peak shapes of these nucleotide phosphates across both methods **(Figure 9C–G)**. Chromatograms show that LT-ZHILIC (green traces) yields sharper and more symmetrical peaks than ZIC-pHILIC (blue traces). However, despite observing clear peaks for all five metabolites from ZIC-pHILIC, our automated analysis pipeline with Compound Discoverer was unable to identify these metabolites. We hypothesize that these nucleotide phosphates were detectable only with LT-ZHILIC because this method provides improved signal-to-noise ratios and substantially narrower elution profiles. For example, ATP exhibited a marked reduction in full peak width—from 3.35 ± 0.09 min with ZIC-pHILIC to 0.81 ± 0.13 min with LT-ZHILIC, supporting this interpretation. Collectively, these results underscore the ability of LT-ZHILIC to deliver deep biological insights through enhanced sensitivity, improved peak shapes, and broader metabolic coverage.

**Figure 9.**
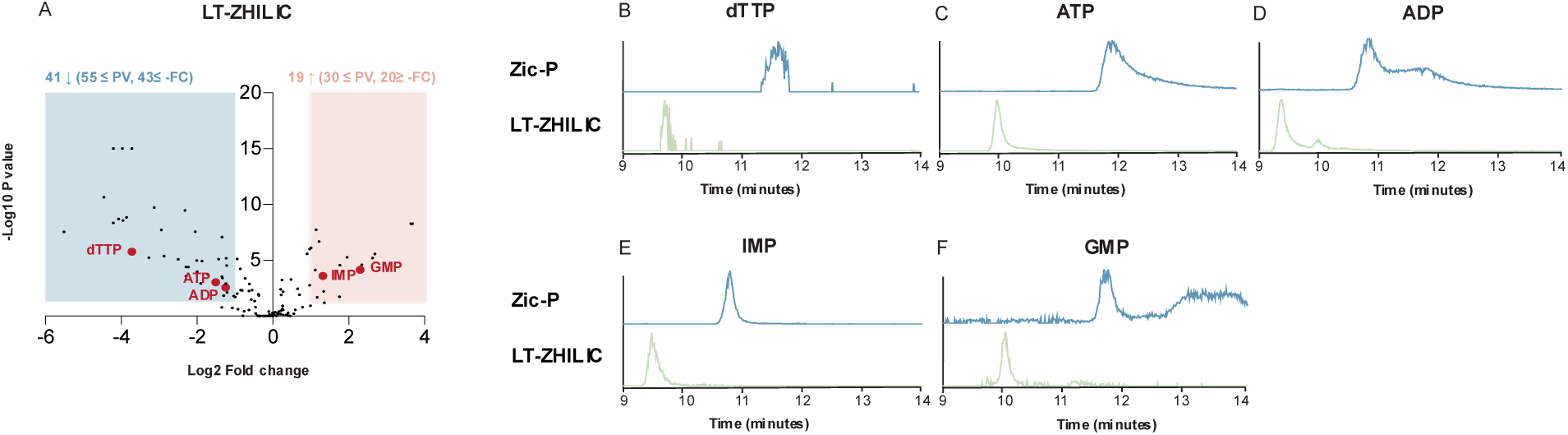
**Nucleotide phosphate species detected in deprivation of glutamine and pyruvate from K562 extractions**. (A–B) Volcano plots of K562 cells cultured under combined glutamine and pyruvate deprivation, analyzed in negative ion mode using ZIC-pHILIC (A) and LT-ZHILIC (B). Unique metabolites are highlighted in red. (C–G) Peak-shape comparison of highlighted metabolites detected using LT-ZHILIC (green) and ZIC-pHILIC (blue).

## Discussion

Since its introduction by Waters^16^, the Z-HILIC column has been applied to metabolomics studies ^24,25^, and is generally used with the ABC B gradient system. However, to our knowledge, no prior reports have systematically described the retention time drift associated with the ABC B system, which can potentially bias quantitative metabolomics results. One previous study observed peak pattern shifts when using standard borosilicate glass bottles and attributed these changes to ion leaching (e.g., sodium, potassium, borate) that altered the mobile phase composition.^26^ Although that study demonstrated improved short-term stability (up to 12 hours) using plastic solvent bottles, our experiments (not shown) demonstrated that this modification did not fully resolve the issue. In our hands, the ABC B mobile phase exhibited progressive retention time drift over four days, consistent with the chemical instability of ammonium bicarbonate. This buffer gradually decomposes into ammonia, carbon dioxide, and water at room temperature, leading to the loss of volatile components, pH drift, and reduced buffering capacity.^27^ Although quaternary pumping systems could, in principle, mitigate some retention challenges by enabling more flexible solvent blending, they are known to exhibit poor reproducibility under the shallow-gradient conditions required for comprehensive metabolome profiling. In mixed aqueous–organic systems, additional factors such as limited solubility and solvent evaporation further accelerate degradation, contributing to signal variability and migration of chromatographic peaks. By contrast, the alternative ACN B method showed no retention drift but yielded poorer chromatographic resolution, highlighting a trade-off between stability and separation performance.

We found substantial improvement to ACN B by operating with the column compartment at 5C. Low-temperature operation, though commonly used in hydrogen-deuterium (HDX) exchange mass spectrometry (MS) to preserve labile features and prevent back-exchange^28^, has rarely been applied in metabolomics, where higher temperatures (typically around or above 30 °C) are favored to improve peak shape and reproducibility.^18,29^ However, we observed that at these higher temperatures, key metabolites such as (iso)citrate, malate, and ATP often produce broad or poorly resolved elution profiles, thereby limiting detection sensitivity and metabolome coverage in untargeted analyses. In addition, redox-active and nucleotide phosphate metabolites are particularly prone to artifactual noise or quantification bias when analyzed using standard methods such as ZIC-pHILIC.^30,31^ By contrast, here we show that LT-ZHILIC produced sharper, more symmetrical peaks for several selected metabolites, resulting in higher quantitative reliability and stability for both targeted and untargeted analyses. While we don’t know the mechanism for this improved separation, we speculate that it may arise from the temperature gradient between room-temperature mobile phases and the chilled column, which can partitioning behavior at the stationary phase interface. At lower temperatures, analytes diffuse more slowly and interact more uniformly with the stationary phase, reducing band broadening and enabling well-resolved peaks for metabolites that elute poorly at conventional temperatures.

Notably, operating at colder temperatures did not result in improved peak shapes for the ZIC-pHILIC column, consistent with a mechanism that may be more specific to the BEH stationary phase chemistry. These results suggest that other columns using BEH-based stationary phases, such as BEH amide columns, could also benefit from lowering column temperature. It is also possible that using a colder column temperature can improve the stability of certain labile metabolites, although this remains speculative and will require future experiments to verify.

Our findings clearly demonstrate that LT-ZHILIC generally outperforms ZIC-pHILIC in resolving biologically essential metabolic pathways, particularly those central to energy metabolism, redox regulation, and nucleotide turnover. The NADH/NAD⁺ ratio is a master regulator of cellular redox homeostasis, governing mitochondrial respiration, oxidative stress signaling, and the activity of numerous metabolic enzymes ^32^. Therefore, the systematic overestimation observed with ZIC-pHILIC introduces a significant risk of misinterpreting redox dynamics. Likewise, nucleotide phosphates such as ATP, ADP, and AMP are foundational to energy transfer, kinase signaling, and metabolic flux; failure to detect ATP in an untargeted workflow, as observed with ZIC-pHILIC can fundamentally distort interpretation of cellular energy states ^33^. In contrast, LT-ZHILIC consistently captures these key metabolites with high fidelity using the same analysis pipeline, providing a markedly more reliable representation of underlying biology. While the mechanistic basis for its enhanced performance warrants further investigation, the evidence presented here establishes LT-ZHILIC as the more robust and biologically accurate platform for metabolomics studies requiring precise quantification of redox and energy-related metabolites.

While the LT-ZHILIC method offers substantial improvements in chromatographic stability and performance, several limitations remain. In our experiments, the cold temperature caused column backpressure to reach approximately 700–1000 bar, which could pose challenges when analyzing more complex matrices or larger sample batches, and thus pressure tolerance should be carefully considered for method scaling. Moreover, operation at low temperatures (≈5 °C) is not universally compatible with all LC systems and may require external temperature-control setups, such as the customized chamber used in our cryogenic experiments. Another area for improvement is the current lack of a dedicated LT-ZHILIC metabolomics spectral library, which would enhance metabolite identification confidence and cross-study comparability. Finally, although the LT-ZHILIC system demonstrated stable retention over four days, longer-term evaluations are needed to fully assess its robustness for extended studies and high-throughput metabolomics workflows. Collectively, our findings indicate that while LT-ZHILIC represents a promising advance, further optimization and resource development are essential for its broader adoption.

## Conclusion

In this study, we characterized two buffer systems for chromatographic LC-MS separation of plasma and intracellular extracts on a Z-HILIC column and found that using high-pH ammonium bicarbonate in buffer B can substantially improve peak shapes, but at the cost retention time drift over the course of four days. We therefore introduce an optimized low-temperature (5 °C) HILIC approach (LT-ZHILIC), that uses only acetonitrile in buffer B and exhibits improved peak shapes and robust retention time stability. Importantly, LT-ZHILIC demonstrated high resolution of targeted analysis of TCA cycle intermediates, redox-sensitive metabolites NADH and NAD^+^, and untargeted analysis of nucleotide phosphates. Compared with the commonly used ZIC-pHILIC column, the LT-ZHILIC method generally performed equivalently or better to minimize variations and enable more reliable quantitative analysis in untargeted metabolomics. Collectively, these findings establish LT-ZHILIC as a robust and versatile chromatographic platform that underscores its potential as a powerful tool for uncovering previously obscured biochemical alterations and guiding future pathway or therapy-focused studies.

## Acknowledgements

We would like to thank Dr. Alexander Ivanov, Purvi Saxena, and Angela Rojas-Merchan for helpful discussions.

## Conflicts of Interest

O.S.S. served as a consultant for Handshake AI. T.K.B. is an employee at Waters.

---

**Supplementary Figure S1.**
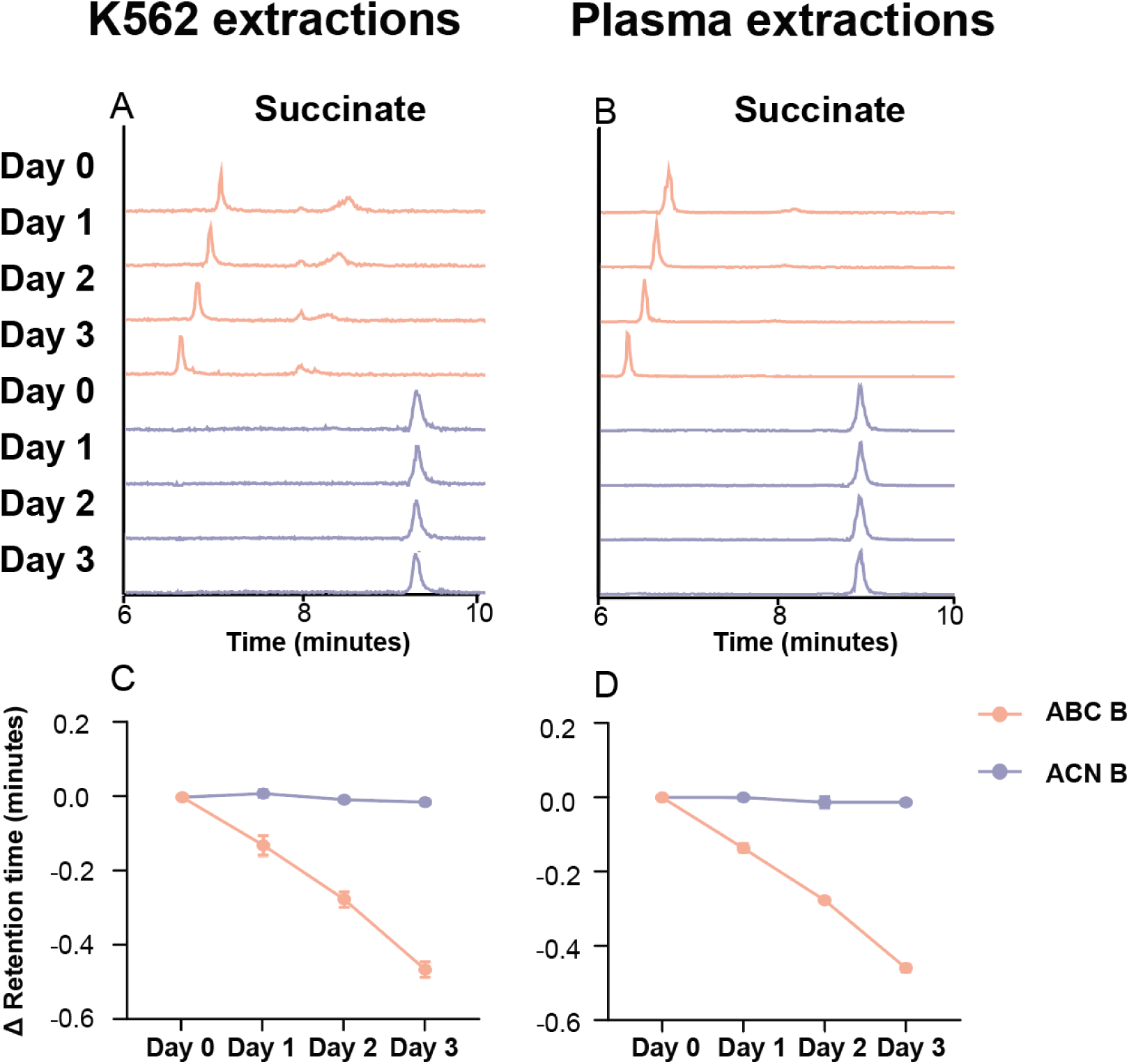
Retention time stability of succinate with ABC B and ACN. **B** (A–B) Representative chromatograms of succinate in K562 cell extracts (A) and plasma samples (B). Metabolites analyzed using the ABC B method are shown in purple; those analyzed using the ACN B method are shown in orange. (C-D) retention time trends for each metabolite over a three-day time course (Day 0 to Day 3) comparing the ABC B (purple) and ACN B (orange) methods.

**Supplementary Figure S2.**
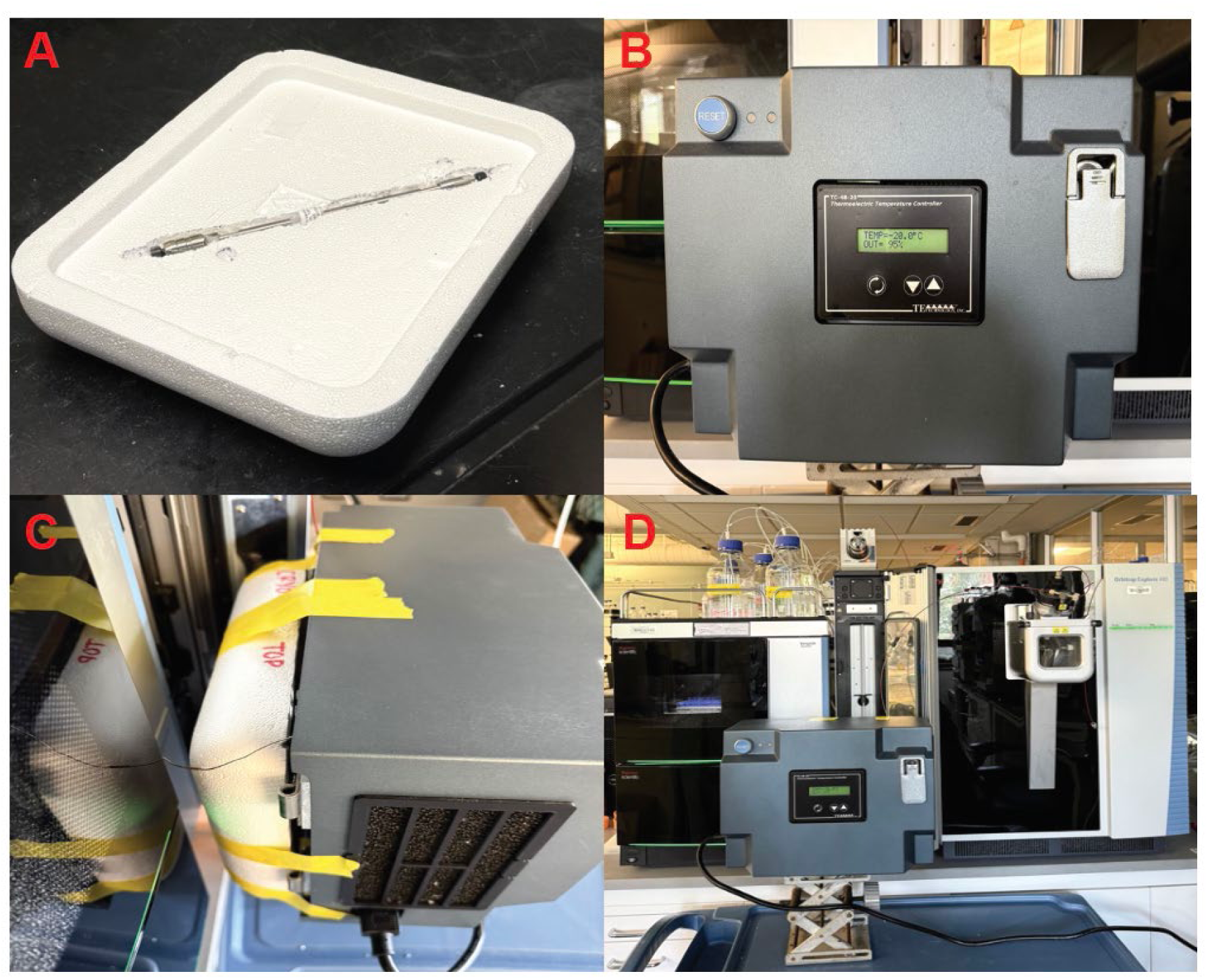
A column compartment setup for cryogenic HILIC. (A–D) Cryogenic setup of the Z-HILIC system. (A) Customized foam insulation designed to fit the Z-HILIC column for temperature preservation. (B) The cryogenic temperature-controlled chamber used for sub-zero operation. (C) The complete cryogenic Z-HILIC setup assembled for testing. (D) Final configuration of the cryogenic Z HILIC system connected to the LC–MS instrument.

**Supplementary Figure S3.**
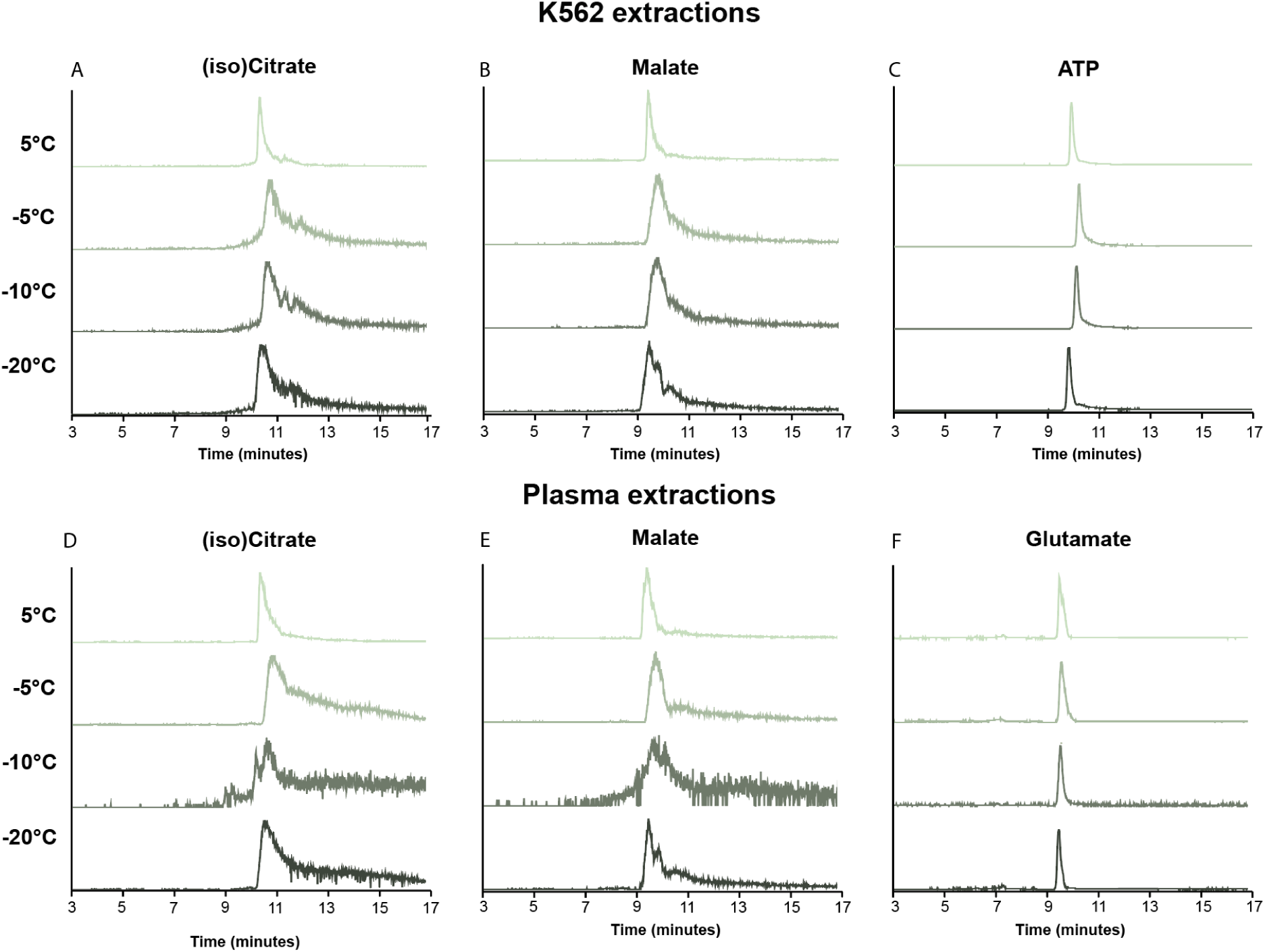
Optimization of the ABC B method under low-temperature conditions (Cryo-HILIC). Selected metabolites were analyzed with progressively lower column compartment temperatures: 5°C, –5°C, –10°C, and –20°C. (A–C) Representative chromatograms of (iso)citrate, malate, and ATP in K562 cell extracts, shown from top to bottom in the order of decreasing temperature. (D–F) (Iso)citrate, malate, and glutamate chromatograms in plasma samples. The optimal peak shape was achieved using a flow rate of 0.15mL/min, a column temperature of 5°C, and an initial mobile phase composition of 80% B.

**Supplementary Figure S4:**
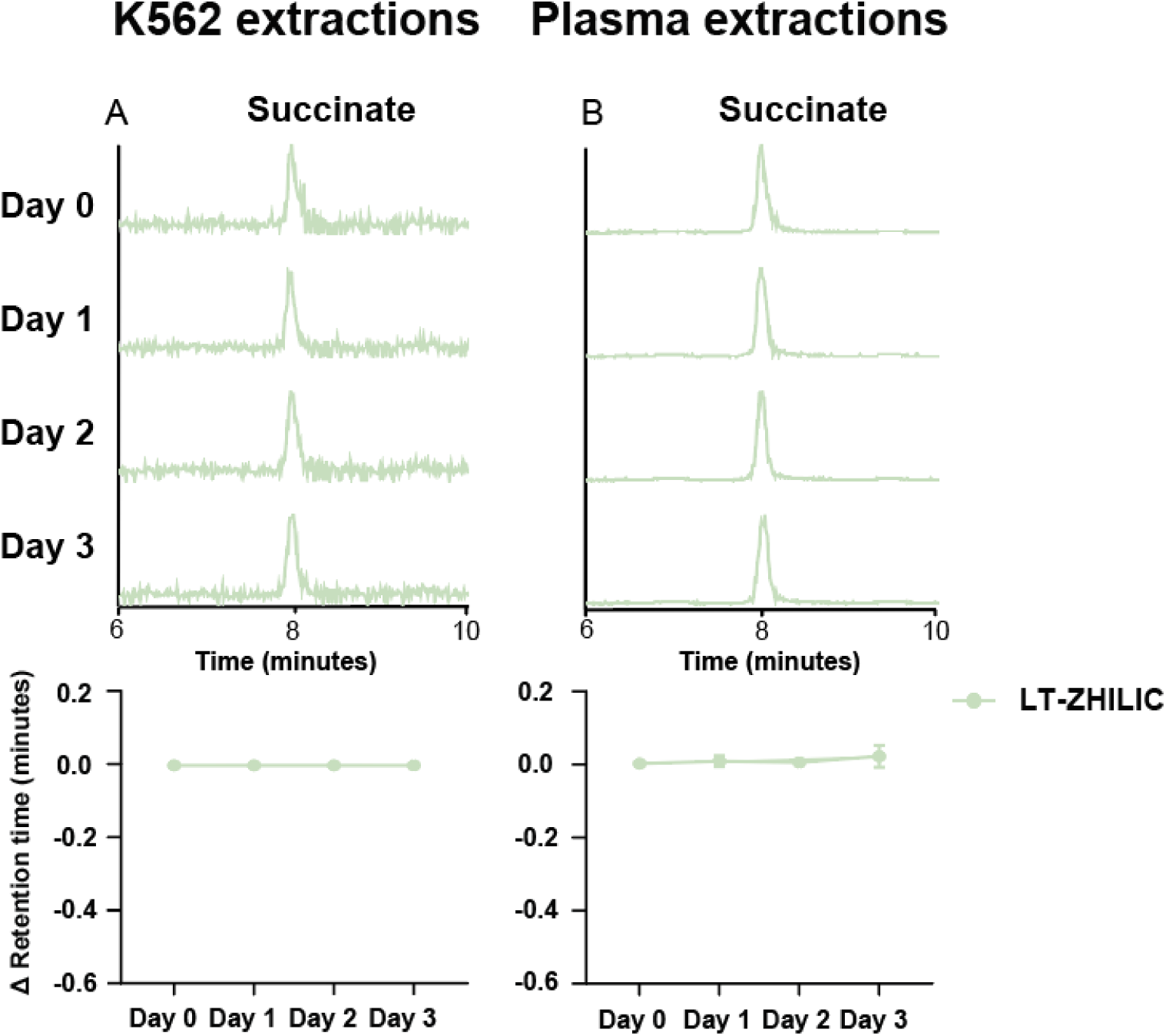
Retention time stability of succinate with LT-ZHILIC. Time-course analysis of LT-ZHILIC stability in K562 and plasma sample extracts. (A–B) Succinate from K562 extracts (A) and plasma extracts (B). Over the four days, LT-ZHILIC demonstrated consistent retention-time stability.

**Supplementary Figure S5.**
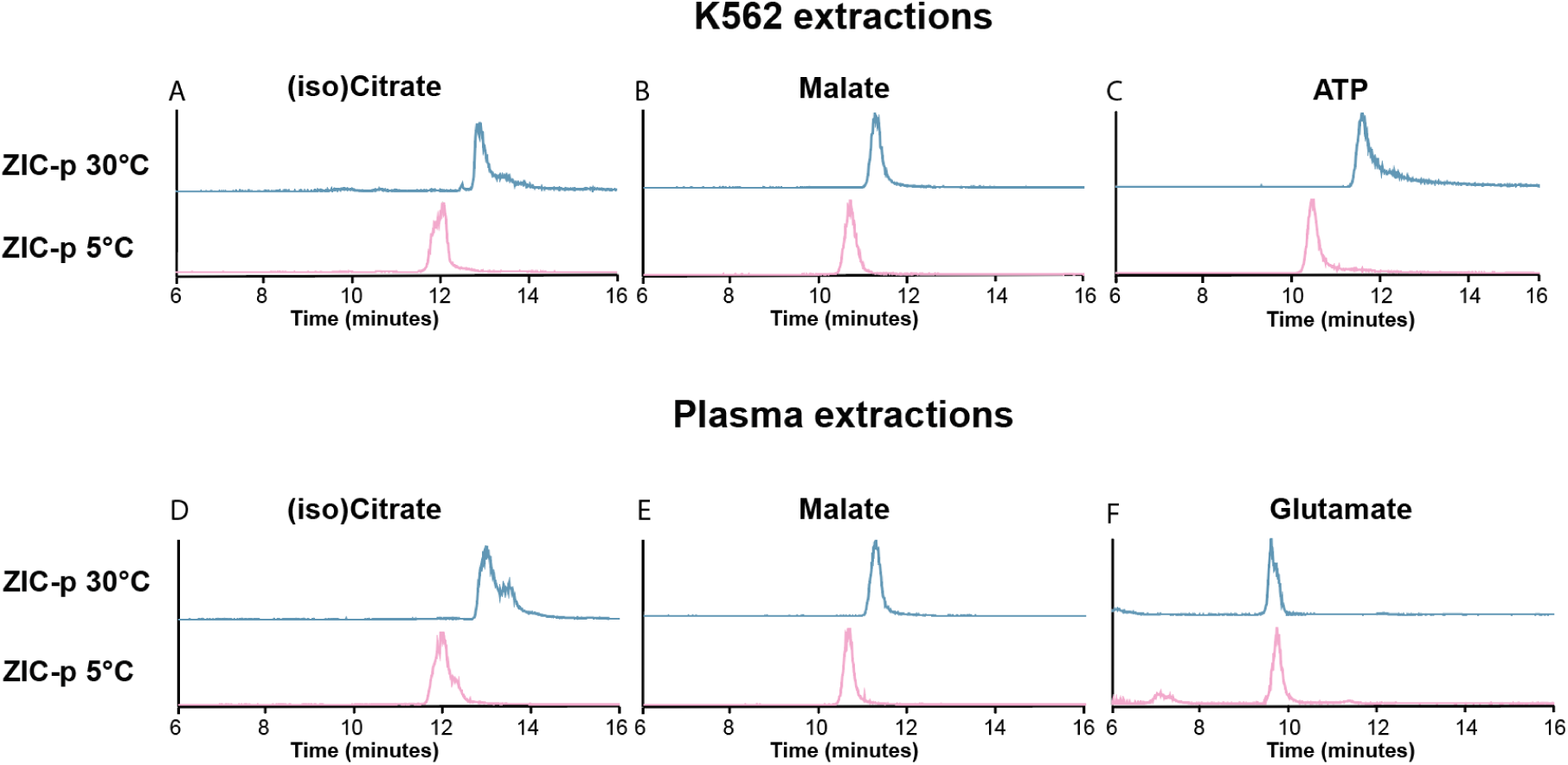
Peak-shape comparison of selected metabolites analyzed on ZIC-pHILIC under different temperatures. Representative results from K562 cell extracts (A-C) and Corresponding results from plasma extracts (D-F). Traces from ZIC-pHILIC at 30 °C are shown in blue, and those at 5 °C in pink.

**Supplementary Figure S6.**
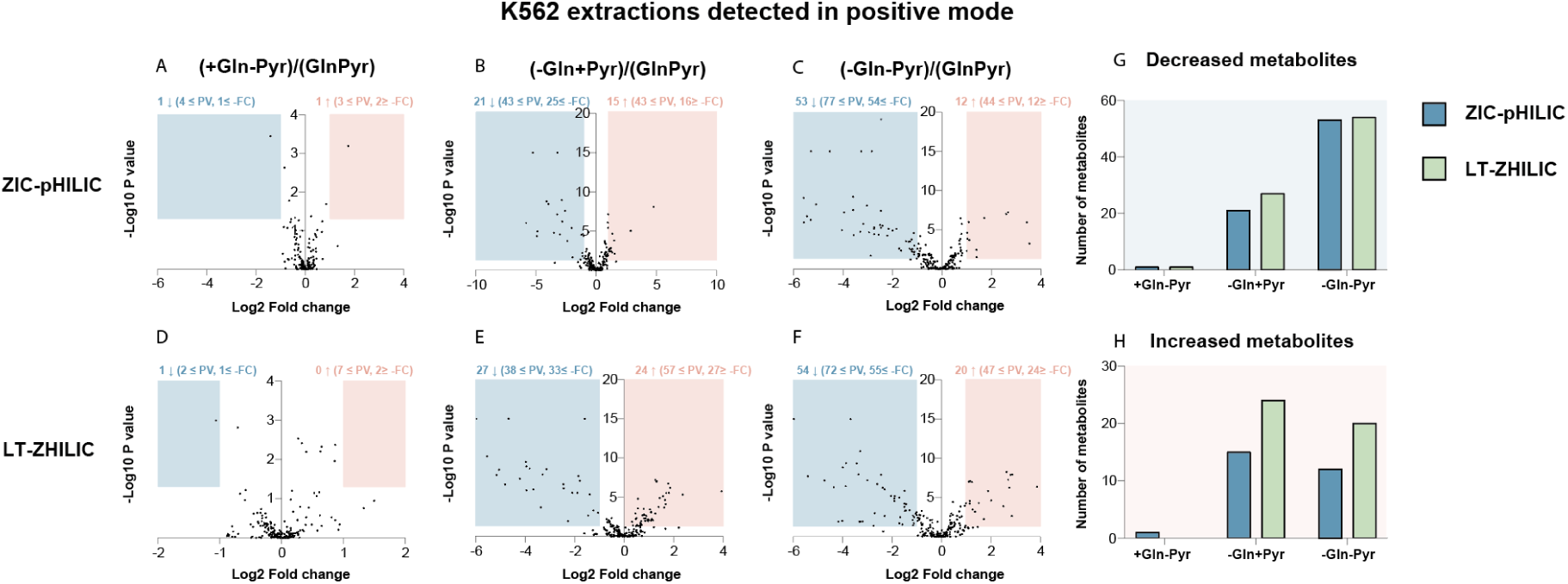
Positive mode volcano plots of K562 cell extracts under different treatment conditions with or without glutamine and pyruvate. Analyzed with Compound Discoverer using data from ZIC-pHILIC (A-C). Corresponding results analyzed using LT-ZHILIC (D-F). (G–H) Summary of downregulated (G) and upregulated (H) metabolites detected using ZIC-pHILIC (blue) and LT-ZHILIC (green).

